# Feeding a rich diet supplemented with the translation inhibitor cycloheximide decreases lifespan and ovary size in Drosophila

**DOI:** 10.1101/2024.08.19.608713

**Authors:** Hye Jin Hwang, Rachel T. Cox

## Abstract

*Drosophila* oogenesis has long been an important model for understanding myriad cellular processes controlling development, RNA biology, and patterning. Flies are easily fed drugs to disrupt various molecular pathways. However, this is often done under poor nutrient conditions that adversely affect oogenesis, thus making analysis challenging. Cycloheximide is a widely used compound that binds to and stalls the ribosome, therefore reducing protein synthesis. Since egg production is a highly nutrient-dependent process, we developed a method to feed female *Drosophila* a rich diet of yeast paste supplemented with cycloheximide to better determine the effect of cycloheximide treatment on oogenesis. We find flies readily consume cycloheximide-supplemented yeast paste. Males and females have reduced lifespans when maintained on cycloheximide, with males exhibiting a dose-dependent decrease. While females did not exhibit decreased egg-laying, their ovaries were smaller and the number of progeny reduced, indicating substandard egg quality. Finally, females fed cycloheximide have disrupted oogenesis, with smaller ovaries, missing ovariole stages, and an increase in apoptotic follicles. Together, these data support that reduced protein synthesis adversely affects oogenesis with a rich diet that provides optimal nutrient conditions. In addition, this method could be used more broadly to test the effect of other drugs on *Drosophila* oogenesis without the confounding effects caused by poor nutrition.

## Introduction

*Drosophila* has been an excellent model system to dissect the genetic pathways and molecular mechanisms controlling processes as diverse as behavior, development, and cell differentiation. Females contain a pair of ovaries made up of ∼20 ovarioles (Fig 1A, B). Each ovariole is a string of developing follicles, with a specialized structure at the anterior called the germarium which houses the germline stem cells (Fig 1C). Each follicle is composed of 16 interconnected cells: one cell will become the oocyte while the other fifteen develop as nurse cells (Fig 1C) (Spradling, 1993). Oogenesis is energy dependent and highly sensitive to nutrition (Armstrong, 2020). Females fed a diet supplemented with wet yeast paste have maximum germline stem cell division and egg production, with all stages of follicle development present (Fig 1C).

**Fig 1.**
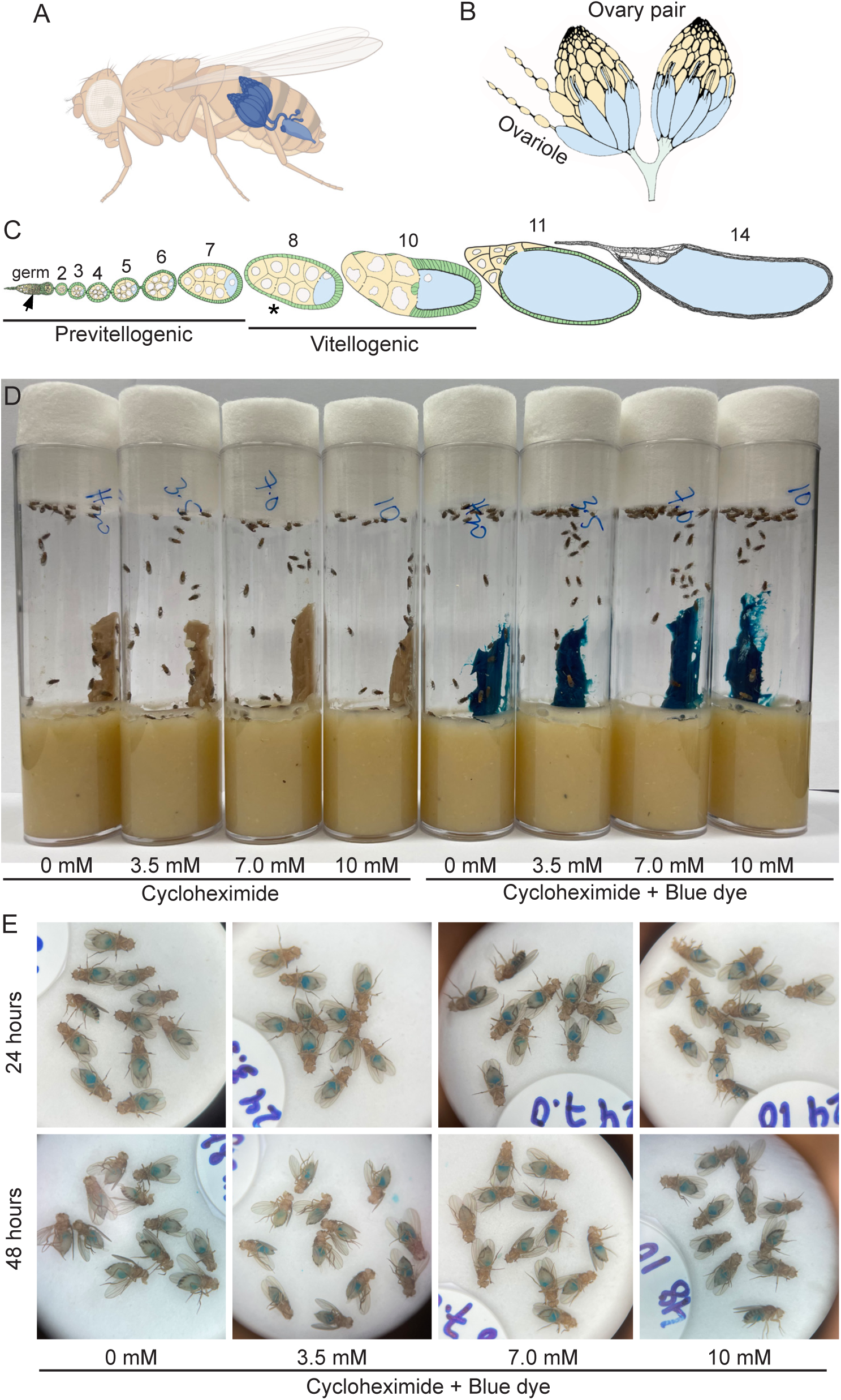
Flies readily consume wet yeast paste supplemented with CHX. (A-C) Schematics of Drosophila female ovaries. (A) Female flies contain a pair of ovaries. (B) Each ovary is composed of strings of developing follicles called ovarioles. (C) Ovarioles in females fed rich diets contain all stages of follicles (Stage 2-14). Stage 8 (asterisk) and Region 2 in the germarium (germ, arrow) are the two developmental stages that undergo apoptosis with nutritional stress. (D) Vials of wild-type flies feeding on wet yeast paste with 0, 3.5, 7.0, and 10 mM CHX. FD&C Blue Dye no. 1 was used to visualize food ingestion by the flies (right side). (E) Blue food is visible in the guts of female flies from the vials in D. Thirty animals were tested for each treatment on the same day.

However, when females are fed a poor diet, oogenesis is quickly affected to conserve energy. A clear indication of nutritional stress is apoptosis that occurs at Stage 8 and missing follicle stages, although there are other problems including mitochondrial mislocalization, changes to metabolism and apoptosis in the germarium (Drummond-Barbosa & Spradling, 2001). This is a conundrum for researchers wishing to disrupt various pathways and study the effect on oogenesis by feeding flies various compounds. The most common method of drug feeding is using a sucrose solution or spiking the standard fly media, neither of which constitutes a rich diet and adversely affects oogenesis (Belozerov et al., 2014; Jeong et al., 2022; Kim et al., 2023; Lagasse et al., 2009; Tully et al., 1994). In addition, a further complication is that wet yeast paste contains active yeast cultures that will presumably also metabolize any compound, thus reducing the concentration of drug.

To circumvent these problems, we developed a method to feed *Drosophila* a rich diet of yeast paste supplemented with cycloheximide (CHX), a translation inhibitor. CHX is a popular compound widely used to study intracellular events and physiological effects by reducing of protein synthesis (Darvishi & Woldemichael, 2016; Jevtov et al., 2015; Kim et al., 2023; Lin et al., 2008; Moutaoufik et al., 2014; Sheth & Parker, 2003). This method would allow for examining how decreased translation and ribosome stalling affects lifespan and oogenesis without the confounding effects of decreased nutrition. Published articles examining the effects of CHX have used the food with limited sources of nutrients when flies were fed with drugs (Kim et al., 2023; Lagasse et al., 2009; Lee et al., 2018; Tully et al., 1994). To support that this feeding method is effective, females fed yeast paste with CHX must readily and reliably consume the drug and exhibit deleterious effects on the fly. Here, we show that feeding CHX in a rich diet reduces lifespan in males and females and females have reduced protein synthesis in the ovary. We also show that feeding CHX using this method does not affect the number of eggs laid, but does affect their quality, resulting in reduced number of progeny. In addition, ovary size is reduced and there is an increased amount of apoptotic and missing follicle stages. Overall, our data supports that CHX treatment directly reduces lifespan and disrupts oogenesis in flies supported with a rich diet. In addition, this method could also be used to feed other drugs to Drosophila in order to test the drugs effect on development and lifespan without the added stress of a poor diet.

## Results

### Flies readily consume wet yeast containing cycloheximide, decreasing protein synthesis

To ensure any CHX effects were due to decreased protein synthesis rather than directly due to poor nutrition that could result from the flies’ food avoidance due to drug taste, we fed flies varying concentrations of CHX mixed in their preferred food source, wet yeast paste. To test whether flies would accept and consume this source of CHX, they were fed freshly made CHX-supplemented yeast paste containing increasing concentrations of CHX for 24 or 48 hrs. We supplemented the food with FD&C blue no. 1 to track food ingestion (Fig 1D). Females readily ate the yeast paste as indicated by the blue guts visible in their abdomens (Fig. 1E). To ensure that CHX was having the desired effect and suppressing protein synthesis, we coupled CHX feeding with puromycin (Fig 2A). Puromycin is a naturally occurring aminonucleoside that mimics tyrosyl-tRNA and incorporates into nascent polypeptides. Since it is not amino acid specific, incorporation terminates all translation. Well-fed flies were fed with CHX-containing fresh yeast paste with CHX for 24 hrs, then transferred to fresh yeast paste containing both CHX and PUR and allowed to feed for an additional 24 hrs (Fig 2A). Fly extract was then probed with anti-puromycin antibodies that recognize the terminated nascent peptides (Fig 2B, upper panels). If CHX decreased translation, we expected a decrease in the amount of puromycin incorporation. Flies fed 3.5, 7.0, and 10 mM CHX had decreased puromycin incorporation, thus supporting that this method of feeding CHX is effective (Fig 2B).

**Fig 2.**
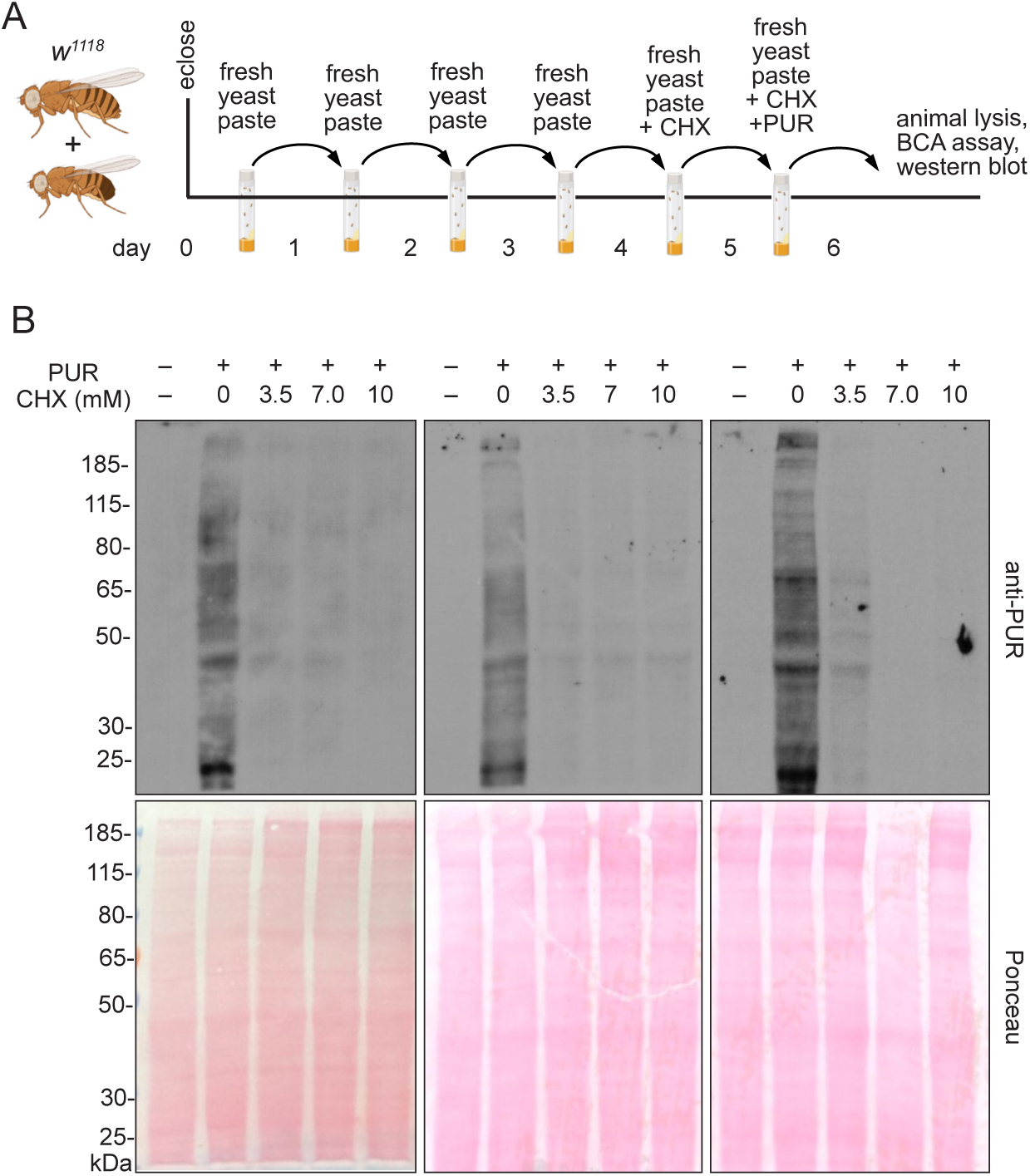
Flies fed CHX-supplemented yeaste paste have reduced protein synthesis. (A) Schematic illustrating the feeding regimen for puromycin (PUR) incorporation in (B). Newly eclosed flies fed freshly prepared yeast paste for four days were given fresh yeast paste containing CHX for 24 hrs, after which they were transferred to the food vial with fresh yeast paste containing both CHX and PUR. Twenty-four hours later, protein was extracted from the female adults and analyzed. (B) Biological triplicate western blots probed with anti-PUR antibodies showing decreased puromycin incorporation with CHX feeding. 40 µg of protein sample was loaded per lane. Ponceau staining (lower blots) are used as a control for gel transfer to the membrane. PUR-: negative control for puromycin feeding.

### Cycloheximide in a rich diet shortens lifespan and depresses larval growth

We next tested the effect of dietary CHX on lifespan (Fig 3). It is possible that CHX treatment would be quickly lethal to flies if our feeding regimen immediately blocked global translation. However, we found the flies tolerated daily dietary CHX when fed freshly made yeast paste supplemented with CHX (Fig 3B, C). We found that continually feeding yeast paste supplemented with 3.5, 7.0, or 10 mM CHX was not immediately lethal but did significantly decrease lifespan in males and females (Fig 3B, C). Males exhibited a dose-dependent decrease in lifespan, whereas females did not (Fig. 3B vs. C). We noticed that the larvae, which hatched from the eggs laid on CHX supplemented yeast paste, appeared small and delayed. To test whether larvae that grew on CHX-supplemented yeast had delayed development, we counted how many pupae developed (Fig 3D). CHX strongly suppressed pupation levels compared to larvae not reared on CHX (Fig. 3E). These data show that constant exposure to CHX in a nutrient-rich diet decreases lifespan and that larvae exposed to CHX struggle to pupate.

**Fig 3.**
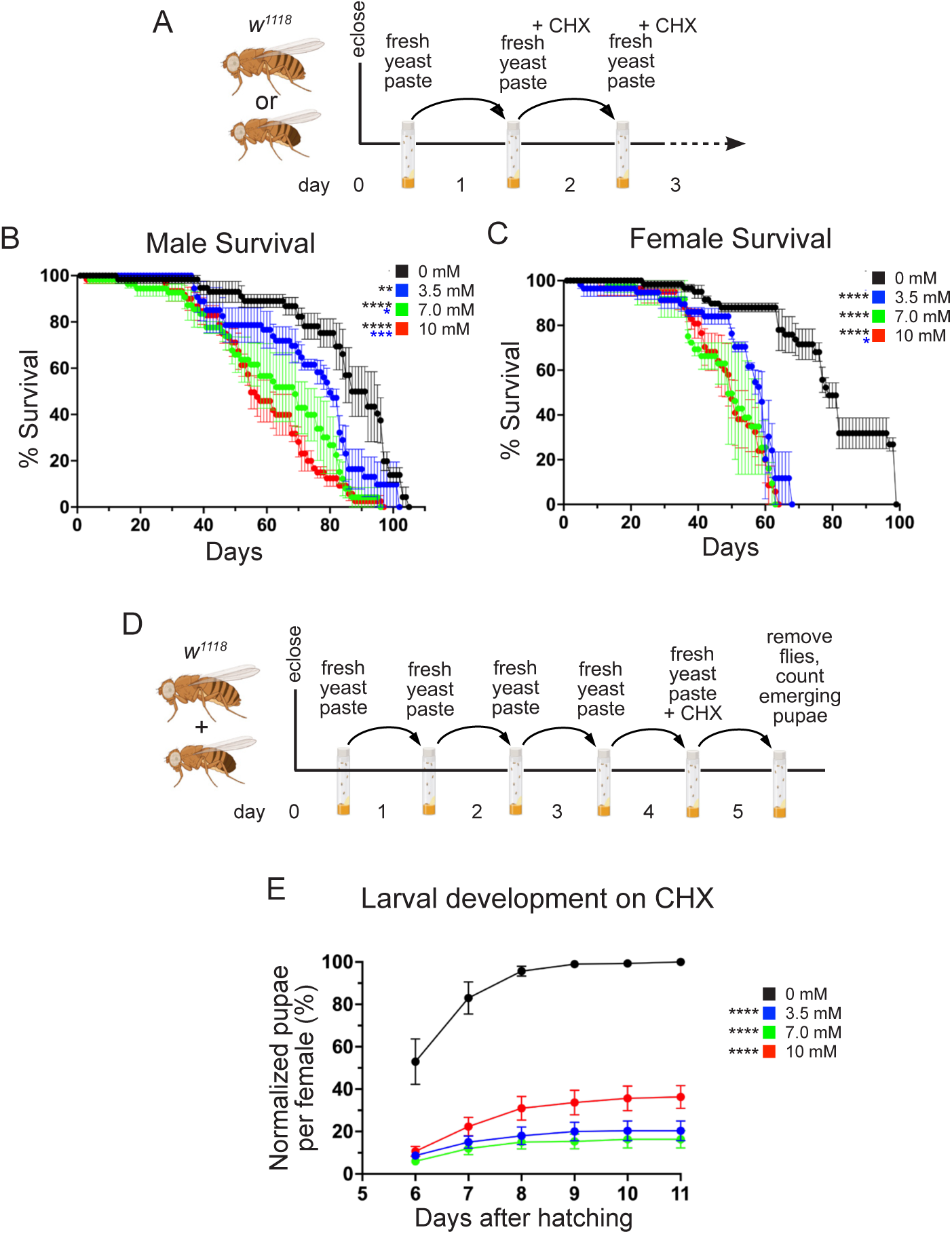
A rich diet supplemented with CHX affects lifespan and larval development. (A) Schematic illustrating the feeding regimen for lifespan measurement in (B, C). Newly eclosed flies were fed freshly prepared yeast paste for one day, after which they were transferred daily to fresh yeast paste containing CHX until animals die. (B, C) Lifespan of flies fed with yeast paste containing varying concentrations of CHX at room temperature. Both males (B) and females (C) have significantly shortened lifespans when fed with CHX compared to controls. Males also show significantly shortened lifespans with increasing CHX concentration. (D) Schematic illustrating the feeding regimen for larval development with CHX in (E). Newly eclosed flies were fed freshly prepared yeast paste for four days, after which they were fed with fresh yeast paste containing CHX for 24 hrs, then removed. The progeny from the eggs laid for 24 hrs were maintained until the pupae emerged. (E) The percent pupae per female fed with varying concentrations of CHX normalized to wild type. CHX-fed progenies have significant growth defects compared to the control group. All experimental conditions had three replicates. (B, C, E) Error bars = standard error (SEM). (B, C) Data were analyzed for significant differences using the Log-Rank test by OASIS2 online application with combined triplicates (B, C), or using ANOVA followed by Tukey’s post hoc test (E). Asterisks in graphs indicate significance as follows: * = p < 0.05, ** = p < 0.01, *** = p < 0.001, ****= p < 0.0001. Asterisks in (E) indicate significance on day 6 after egg laying. The color of asterisks in the graphs denotes the concentration of comparison.

### Females fed a rich diet with CHX have reduced egg quality, but not quantity

Once we had established the feeding flies a rich diet with CHX was having an effect, we wanted to determine how decreased protein synthesis affects oogenesis with optimal nutrition. We found that egg laying was not affected, even at the highest dose of CHX (10 mM) (Fig 4A, B). To test the quality of the eggs laid, we again performed a pupation assay but transferred the CHX-fed females to a standard food vial, letting these females lay eggs in the vial without CHX. (Fig. 4C). With this assay, fewer larvae pupated from females fed CHX in yeast paste, compared to control, with a dose dependent effect (Fig. 4D). These data suggest that, while the number of eggs being laid is unaffected with dietary CHX, egg quality is reduced.

**Fig 4.**
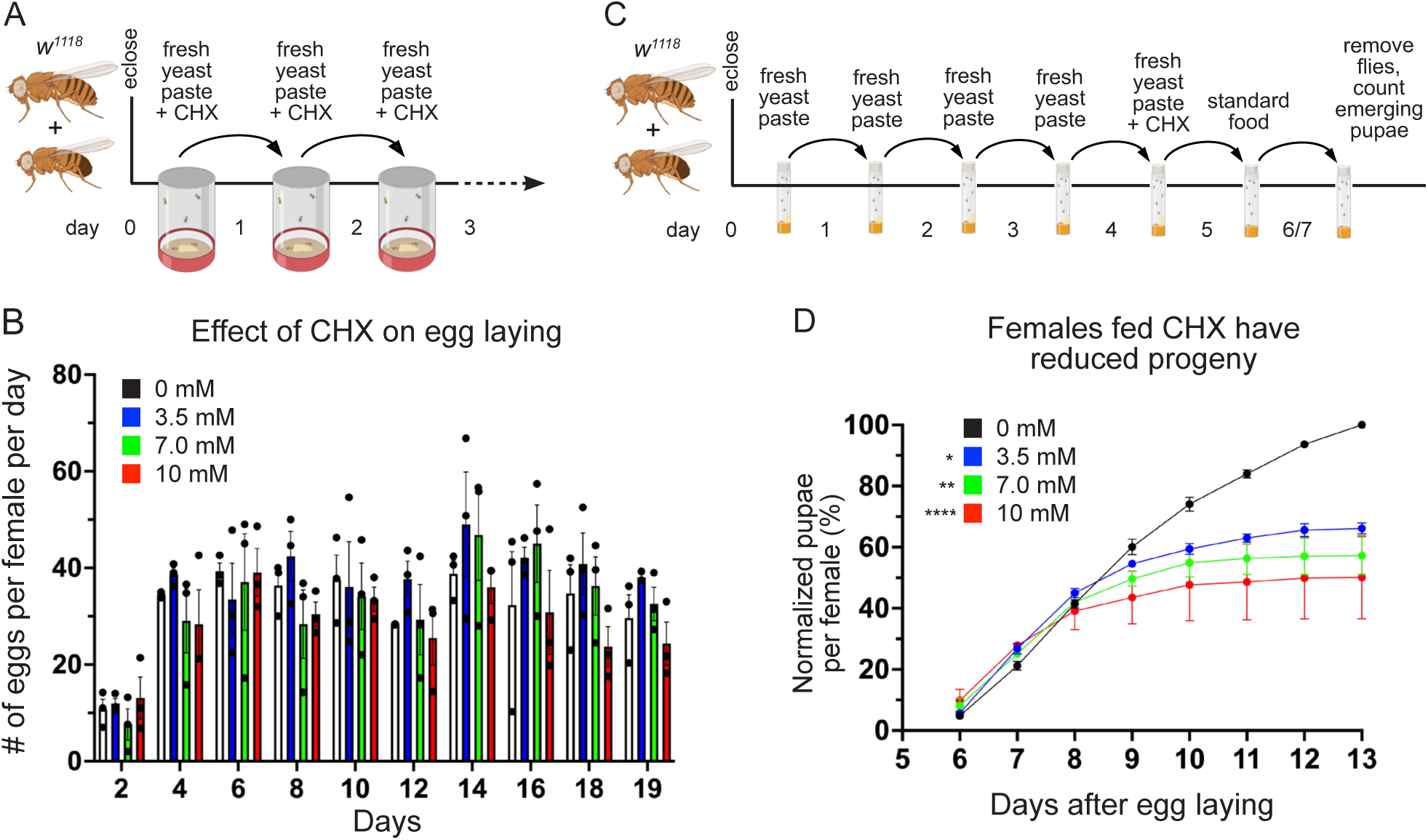
A rich diet supplemented with CHX affects egg quality but not quantity. (A) Schematic illustrating the feeding regimen for egg laying measurement in (B). Newly eclosed flies were fed freshly prepared yeast paste for one day, after which they were transferred daily to an egg-laying cup with fresh yeast paste containing CHX. (B) The number of eggs laid per female fed with varying concentrations of CHX. The total number of eggs was counted every two days. (C) Schematic illustrating the feeding regimen for progeny developmental assays in (D). Newly eclosed flies were fed freshly prepared yeast paste daily for four days, after which they were fed fresh yeast paste containing CHX for 24 hrs. Flies were transferred to a standard food vial and maintained for 48 hrs, after which they were removed. The progenies from the eggs laid for 48 hrs were maintained until pupae emerged. (D) The normalized percent pupae per female fed with varying concentrations of CHX for 24 hrs. The number of pupae from flies fed with CHX-containing yeast paste was significantly decreased compared to control. All experimental conditions had three replicates. (B) Points on the graph represent each of three replicates. Bars on the graph represent the average of all three replicates. (B, D) Error bars = SEM. Data were analyzed for significant differences by ANOVA followed by Tukey’s post hoc test. Asterisks in graphs indicate significance on day 11 after egg laying. * = p < 0.05, ** = p < 0.01, ****= p < 0.0001. The color of asterisks in the graphs denotes the concentration of comparison.

### Females fed a rich CHX diet have ovary defects

Although egg-laying levels remained the same, since the progeny number was reduced, it was possible that there were defects during oogenesis. To identify ovary defects, well-fed females were fed CHX-supplemented yeast paste for 24 hours before ovary dissection (Fig 5A). Whole ovaries dissected from females fed 3.5, 7.0, and 10 mM CHX did not look grossly abnormal but did look slightly smaller compared to control (Fig 5B-E). Females fed CHX for 48 hours had noticeably smaller ovaries (Fig S2). To determine any decrease in ovary size after 24 hours of feeding, we measured the area and perimeter of each ovary (Fig 5F, G). Females fed all three doses of CHX had decreased ovary size, with 10 mM CHX causing the smallest ovaries (Fig 5F, G). Oogenesis is metabolically and energy-intensive and the smaller ovaries might have been caused by less material being made due to reduced protein synthesis. Thus, we extracted and measured total protein from ovaries exposed to CHX. Normalizing for size differences using ovary area, we did not observe a decrease in total protein after 24 hrs of dietary CHX (Fig S2G). To determine whether a rich CHX diet altered follicle stage development, we used indirect immunofluorescence. In well-fed females, all stages (2-14) of follicle development in the ovariole are present (Fig 1C). There are two developmental checkpoints during oogenesis when follicle death occurs: in region 2B of the germarium and Stage 8 (Fig 1C, arrow, asterisk, respectively). This occurs in response to several stressors, of which the best characterized is nutritional stress (Drummond-Barbosa & Spradling, 2001). To identify any missing follicle stages, we labeled ovaries with the antibody against the germplasm protein Vasa and DAPI to label nuclei. Well-fed females had ovarioles that contained the expected follicle stages with no Stage 8 follicles containing condensed nuclei (Fig 6A-A”, Fig S1A-A”). In contrast, females fed a rich diet supplemented with CHX were frequently missing stages and had increased numbers of condensed nuclei, signifying apoptotic follicles (Fig 6B’, B”, Fig S1B-D”). Increasing the dose of CHX increased the number of defective ovarioles (Fig 6C, Fig S1). In the follicles present, we did not detect changes to the number of germ cells per cyst, nor an increase in the loss of the oocyte (Fig S1). These observations support that, with a rich diet, CHX decreases ovary size by increasing apoptotic follicles.

**Fig. 5.**
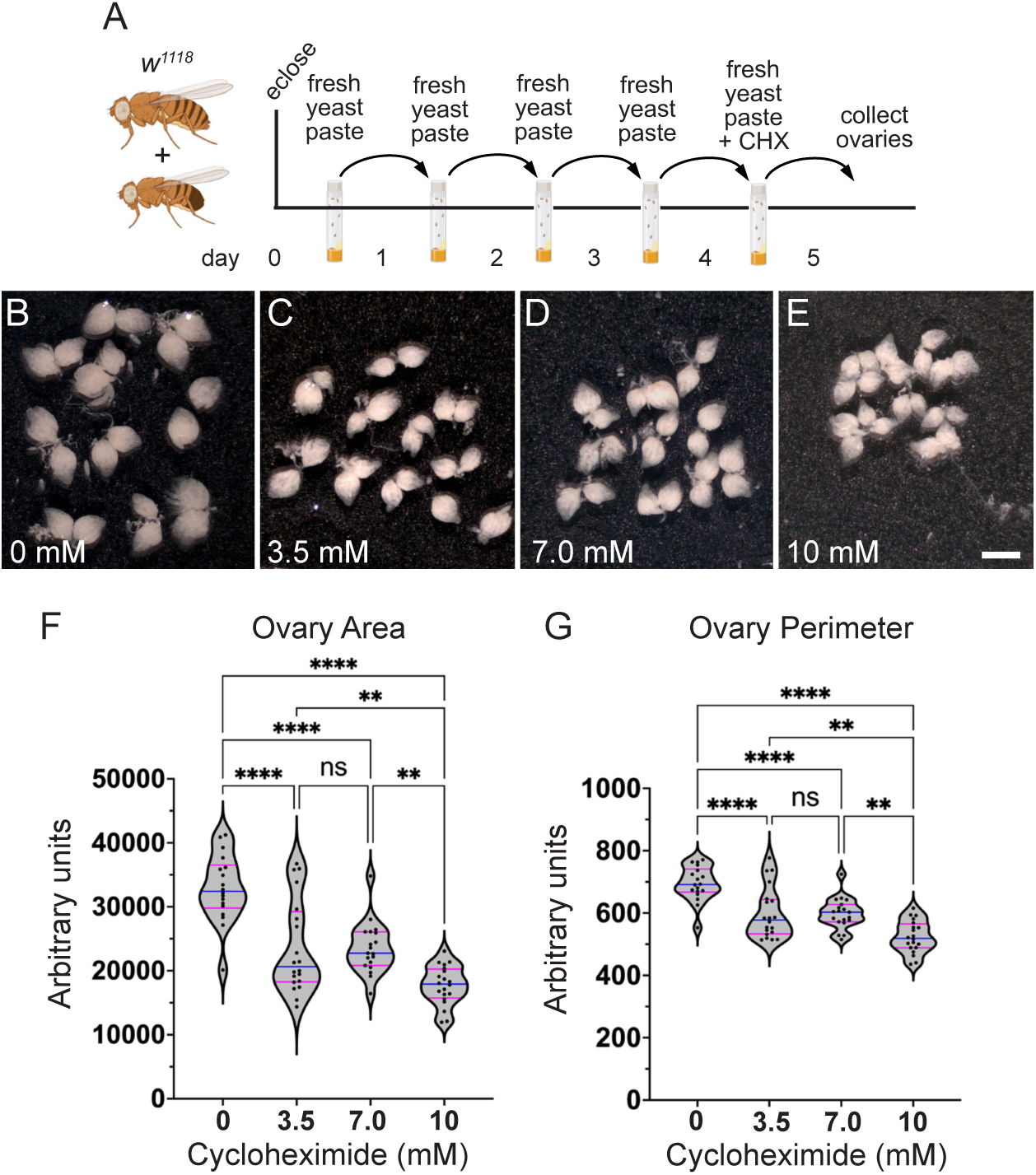
A rich diet supplemented with CHX affects ovary size. (A) Schematic illustrating the feeding regimen for measuring ovary size in (B-G). Newly eclosed flies were fed freshly prepared yeast paste daily for four days, then fed with fresh yeast paste containing CHX for 24 hrs after which the ovaries were dissected. (B-E) Ten pairs of ovaries from the females fed with varying concentrations of CHX for 24 hrs. Females fed CHX have smaller ovaries. The pictures shown are representative of triplicates. These ovaries were analyzed using FIJI/Image J for the whole ovary area (F) and the perimeter (G). (F, G) Violin plots for ovary area (F) and perimeter (G) from females fed CHX. Each point on the violin plot represents one ovary. The median and quartiles are represented by blue and magenta horizontal solid lines, respectively. The plots shown are representative of triplicates. The results shown are combined with triplicates. Error bars = SEM. Data were analyzed for significant differences by ANOVA followed by Tukey’s post hoc test. Asterisks in graphs indicates significance as follows: ** = p < 0.01, ****= p < 0.0001, ns= no significance. Scale bar = 1 mm in E for B-E.

**Fig. 6.**
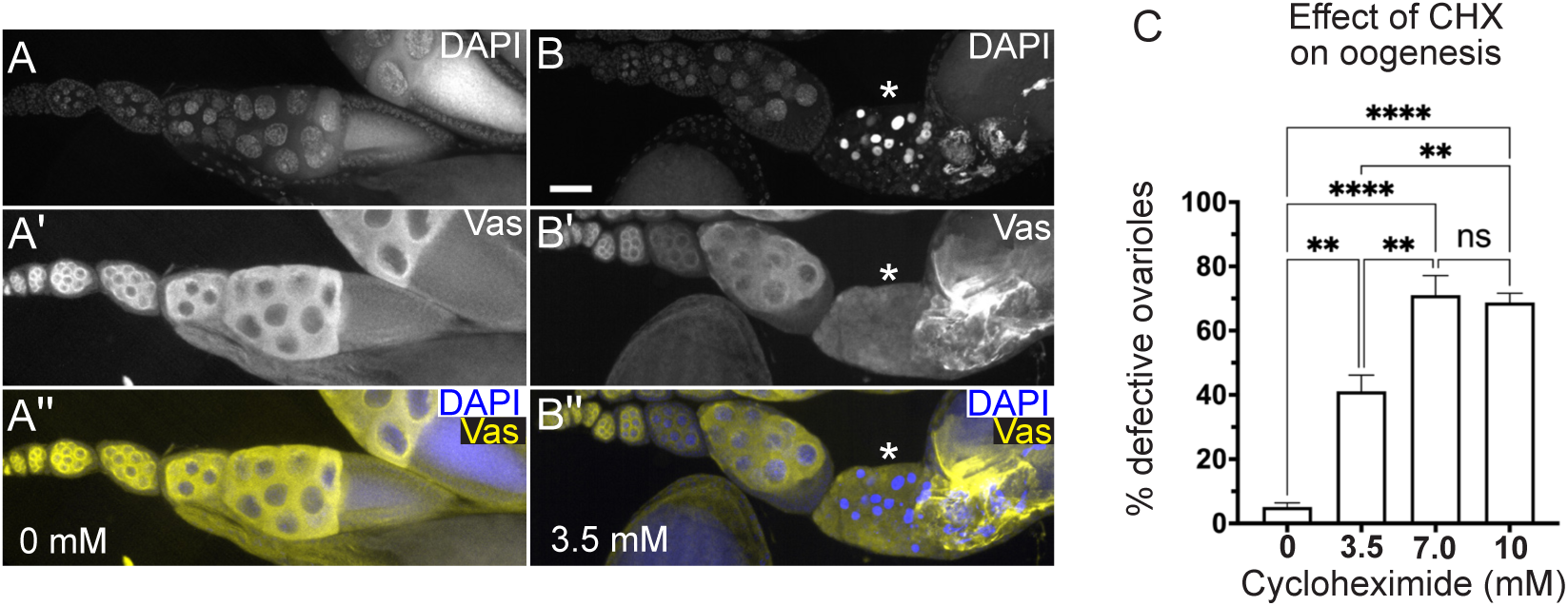
A rich diet supplemented with CHX affects oogenesis. (A-C”) Ovarioles labeled with anti-Vasa antibodies to label germplasm and DAPI to label nuclei. (A-A”) Representative ovariole dissected from a well-fed females show a normally developed ovariole. (B-B”) Dissected ovariole from a female fed with 3.5 mM CHX for 24 hrs. Ovarioles from CHX fed females have an increased number of mid-stage apoptotic follicles as indicated by condensed DAPI-labeled nuclei (B, B”, asterisks). (C) The relative number of ovarioles having developmental defects shown in (A-B”, Fig SF1). Ovarioles with developmental defects contained follicles with apoptotic nuclei (asterisk in B-B”) and/or missing follicle stages (Fig SF1). Approximately fifty to ninety ovarioles were counted in an individual experimental condition (see Materials and Methods for details.) Error bars = SEM. Data was combined with three independent experiments and analyzed for significant differences by ANOVA followed by Tukey’s post hoc test. Asterisks in the graph indicate significance as follows: ** = p < 0.01, ****= p < 0.0001 and ns = not significant. (A, B) White = DAPI, (A’, B’) white = anti-Vasa, (A”, B”, merge) yellow = anti-Vasa, blue = DAPI. Scale bar = 50 µm in B for A-B”.

## Discussion

Dietary CHX has been administered to flies in several contexts, but not for its effects on the *Drosophila* ovary and not by feeding a rich diet (Cervantes-Sandoval et al., 2016; Jelen et al., 2023; Jeong et al., 2022; Lagasse et al., 2009; Lee et al., 2018; Takakura et al., 2021; Tully et al., 1994). A previous study examined the effects of CHX feeding on various developmental stages and found a dose-dependent effect on lifespan (Marcos et al., 1982). Behavioral studies have found that CHX feeding affects certain types of memory in flies (Jelen et al., 2023; Lagasse et al., 2009; Lee et al., 2018; Takakura et al., 2021; Tully et al., 1994). These feeding studies used a standard method that relies on a mixture of sucrose and desired drug and did not use a rich diet, which is critical for supporting optimal ovary development (Drummond-Barbosa & Spradling, 2001; Wei & Lilly, 2014). A sugar-only diet is considered a poor food source for females and does not support robust ovary development and egg laying. Since we wanted to observe the effects of CHX on oogenesis without the confounding effects of a poor diet, we fed flies CHX mixed in live yeast paste, a rich diet that *Drosophila* prefer. Since the yeast in the paste is active and likely metabolizing the CHX, the paste was made fresh daily. This ensured that CHX levels could be maintained at as steady a level as possible, although there was likely a drop in concentration over the 24 hours. Using this method, we saw reduced survival, with females more affected than males. Males, however, exhibited a a dose-dependent lifespan decrease, whereas females did not. Somewhat counterintuitively, female egg-laying was not affected. Given that there was increased loss of follicle stages, it is possible that the rich diet combined with stalled translation increased germline stem cell division, making up for any deficit in ovariole development, but we did not test germline stem cell division rates directly. While egg laying was normal, progeny number was reduced from females fed CHX indicating that the eggs laid were less competent to develop. Dietary CHX reduced ovary size likely due to increased apoptosis at a well-known checkpoint but normalized total protein levels were unchanged. Thus, any action of CHX on disrupting follicle developing appears to be independent of decreased global protein. Rather, these defects could be due to reduced translation of specific transcripts or signaling pathways while the robust signaling pathways responding to a rich food source remain intact. Overall, these data demonstrate that CHX treatment and stalled ribosomes specifically trigger follicle loss and increase apoptosis. In addition, this feeding method could be used for other compounds to distinguish specific effects on ovary development, as opposed to decreased nutrition.

Competing interests: No competing interests declared.

## Materials and Methods

### Fly stocks

Fly stock *w^1118^* was maintained on standard cornmeal fly media. Animals were grown at room temperature. A newly eclosed fly is considered day 0.

### Preparation of cycloheximide solution for wet yeast paste

Cycloheximide (CHX, Sigma-Aldrich, Burlington, MA, USA, cat#. C7698) was dissolved in water to prepare a 10 mM stock solution that was aliquoted and stored at –20 °C until use. The desired concentration of CHX was prepared by serial dilution of the stock solution. To make fresh yeast paste containing CHX, 0.3 g of active dry yeast (Red Star® Yeast) was mixed with 450 µL of CHX solution to create a yeast paste, which was provided daily as needed.

### Blue dye feeding with cycloheximide

Blue-dye feeding was performed as described in Wong et al (Wong et al., 2009). Newly eclosed flies were fattened with yeast paste for 3-7 days. Thirty female flies were then transferred to a vial with fresh fly food containing wet yeast paste (control) or freshly made yeast paste with varying concentrations of CHX prepared with 2.5% (w/v) of FD&C Blue Dye no. 1 (Sigma-Aldrich, Burlington, MA, USA, cat#. 80717).

### Cycloheximide feeding for survival assay

Twenty female or male day 0 adult flies were collected in standard fly food vials containing fresh wet yeast paste. After 24 hrs, the flies were transferred to freshly made wet yeast paste containing CHX. Fresh yeast paste with CHX was provided daily until no flies survived. Flies were maintained at room temperature, and the dead and missing animals were counted every day.

All individual experiments had three replicates. Survival percentage was plotted using GraphPad Prism (GraphPad Software, Boston, Massachusetts USA, www.graphpad.com), and the data was analyzed with the Kaplan-Meier test and for significant differences using the Log-Rank test by OASIS2 online application with combined triplicates (Han et al., 2016).

### Cycloheximide feeding for egg-laying assay

Five newly eclosed female flies and three newly eclosed male flies were collected and fed with wet yeast paste for 24 hrs in an egg-laying cup containing an agar-corn syrup plate. The flies were transferred to a plate supplemented with wet yeast paste supplemented with varying concentrations of CHX. Every 24 hrs, the plate was replaced with fresh CHX-wet yeast paste. The eggs laid on the plate were counted every other day. Each experimental condition had three replicates. The bar graph was generated using GraphPad Prism.

### Cycloheximide feeding for pupation assay

Ten female day 0 adult flies and five male day 0 adult flies were collected in a standard food vial with wet yeast paste. Flies were fed yeast paste for four days. On day 4, flies were transferred to a standard food vial with wet yeast paste containing CHX. In 24 hrs or 48 hrs after CHX feeding, flies were transferred to a new standard food vial that was kept at room temperature. After 48 hrs, flies were removed, and the vials were kept at room temperature. The resulting pupae was counted daily and normalized with the number of parent females. All experimental conditions had three biological replicates. The line graph was generated using GraphPad Prism, and the data was analyzed statistically with ANOVA with multiple comparisons followed by Tukey’s post hoc test.

### Cycloheximide feeding for ovary analysis

Ten female day 0 adult flies and ten male day 0 adult flies were collected in a standard food vial with wet yeast paste. The food vial with fresh wet yeast paste was switched every day for four days. On day 4, all female flies in each vial were transferred to a standard food vial containing freshly made yeast paste with varying concentrations of CHX. Fly ovaries were dissected in Grace’s insect media 24 hrs after CHX feeding. To measure whole ovary size, the dissected ovaries were immediately imaged under a stereomicroscope using an Accu-scope 3076 digital microscope 0.67x-4.5x (Accu-Scope, Commack, NY, USA). After imaging, ovaries were washed with phosphate-buffered saline twice, frozen, and kept at –80 °C for protein analysis. Ovary area and perimeter were measured using Fiji ImageJ (Schindelin et al., 2012). The violin plot was generated using GraphPad Prism, and the data was analyzed statistically using ANOVA with multiple comparisons followed by Tukey’s post hoc test. To evaluate CHX effect on oogenesis, the ovaries were immunostained for Vasa and DNA and imaged as described below. Ovarioles were considered having developmental defects if they contained an apoptotic follicle and/or were missing stage 8-11 egg chambers. The following ovarioles were counted per individual experimental condition: Experiment 1: 78 (0 mM CHX), 92 (3.5 mM CHX), 97 (7 mM CHX), 72 (10 mM); Experiment 2: 44 (0 mM CHX), 72 (3.5 mM CHX), 89 (7 mM CHX), 84 (10 mM CHX); Experiment 3: 68 (0 mM CHX), 81 (3.5 mM CHX), 59 (7 mM CHX), 84 (10 mM CHX). The relative percentage of defective ovarioles compared to normal ovarioles was calculated within each experimental condition. The bar graph was generated by combining the results from three independent experiments using GraphPad Prism, and the data was analyzed statistically using ANOVA with multiple comparisons followed by Tukey’s post hoc test. All individual experiments had three biological replicates.

### Puromycin incorporation assay

Fifteen day 0 adult flies and fifteen male day 0 adult flies were collected in a standard food vial with wet yeast paste. The food vial with fresh wet yeast paste was switched every day for four days. On day 4, all adult flies in each vial were transferred to a standard food vial containing a wet yeast paste with varying CHX concentration. After twenty-four hours, flies were transferred to a standard vial containing a wet yeast paste with both 600 µM puromycin (PUR, Gibco^TM^, Waltham, MA, USA, cat#. A1113803) and varying CHX concentrations. After feeding PUR and CHX, the female flies were collected and kept in –80°C until preparing and quantifying protein amounts as described below. 40µg of each sample was separated by SurePAGE™, Bis-Tris gel (GensScript, Piscataway, NJ, cat#. M00653) and transferred to a nitrocellulosemembrane. The nitrocellulose membrane was probed with mouse anti-puromycin [3RH11] antibody (Kerafest, Shirley, MA, cat#. EQ0001) following a staining with Ponceau S solution (Sigma-Aldrich, Burlington, MA, USA, cat#. P7170) for total protein loading imaging.

### Preparation and quantification of cellular proteins

Dissected and frozen ovaries or frozen whole animals were homogenized with a whole cell lysis buffer composed of 50 mM Tris (pH 8.0), 150 mM sodium chloride, 1 mM ethylenediaminetetraacetic acid (EDTA), 1% NP-40, 0.1% sodium dodecyl sulfate, and Complete-mini EDTA-free protease inhibitor (Roche Applied Science, Indianapolis, IN, USA, cat#. 11836170001) by repeated freeze-thaw cycles and 30 times of strokes with a pestle. Insoluble material was removed by centrifugation (12,000 g for 20 mins at 4 °C), and the subsequent supernatant was collected as a fraction of whole cellular proteins. Protein concentration was determined using Pierce BCA Protein Assay Kit (ThermoFisher Scientific, Waltham, MA, USA cat#. 23227).

### Immunostaining

Ovaries were dissected with Grace’s Insect Medium (Invitrogen, Waltham, MA, USA, cat#11595030) and fixed for 20 mins (4% paraformaldehyde and 20 mM formic acid in Grace’s Insect Medium). After washing with Antibody wash (0.1% Triton X-100 and 1% BSA in phosphate-buffered saline) three times for 20 mins, the tissue was stained with primary antibody rat anti-vasa (1:14, Developmental Studies Hybridoma Bank, Iowa City, IA, USA), overnight at 4 °C. After washing with Antibody wash three times for 20 mins, the tissues were stained with secondary antibody goat anti-rat IgM Cross-Adsorbed Secondary Antibody, DyLight™ 488 (1:500, Invitrogen, Waltham, MA, USA, cat#. SA510010), overnight at 4 °C, then incubated with Antibody wash twice for 20 mins and stained with 5 ng/mL 4’,6-Diamidino-2-phenylindole (DAPI) solution for 10 mins. After removing the DAPI solution, the tissues were mounted in Vectashield Antifade Mounting Medium (Vector Laboratories, Newark, CA, USA, cat#. H-1000). Images were obtained using a Zeiss LSM 980 confocal laser scanning microscope at Plan Apochromat 10x objective (Carl Zeiss Microscopy LLC, White Plains, NY, USA) or a Nikon Eclipse Ti2 spinning disk microscope at Plan Apo λ 10x objective (Nikon Corporation, Tokyo, Japan) with z-stacked for a depth of 20 µm.

## Acknowledgements

We would like to thank the USUHS Biomedical Instrumentation Core and Dr. Dennis McDaniel for imaging support. Antibodies obtained from The Developmental Studies Hybridoma Bank, created by the NICHD of the NIH and maintained at The University of Iowa, Department of Biology were used in this study. Stocks obtained from the Bloomington Drosophila Stock Center (NIH P40OD018537) were used in this study. Cartoons in Figures 1-5, S2 are courtesy of BioRender.

## Funding

This work was supported by the National Institutes of Health GM127938 to R. T. C. The content is solely the responsibility of the authors and does not necessarily represent the official views of the National Institutes of Health, the Department of Defense, or Uniformed Services University.

## Data availability

All relevant data can be found within the article and its supplementary information

## Figure Legends

**Supplementary Fig 1.**
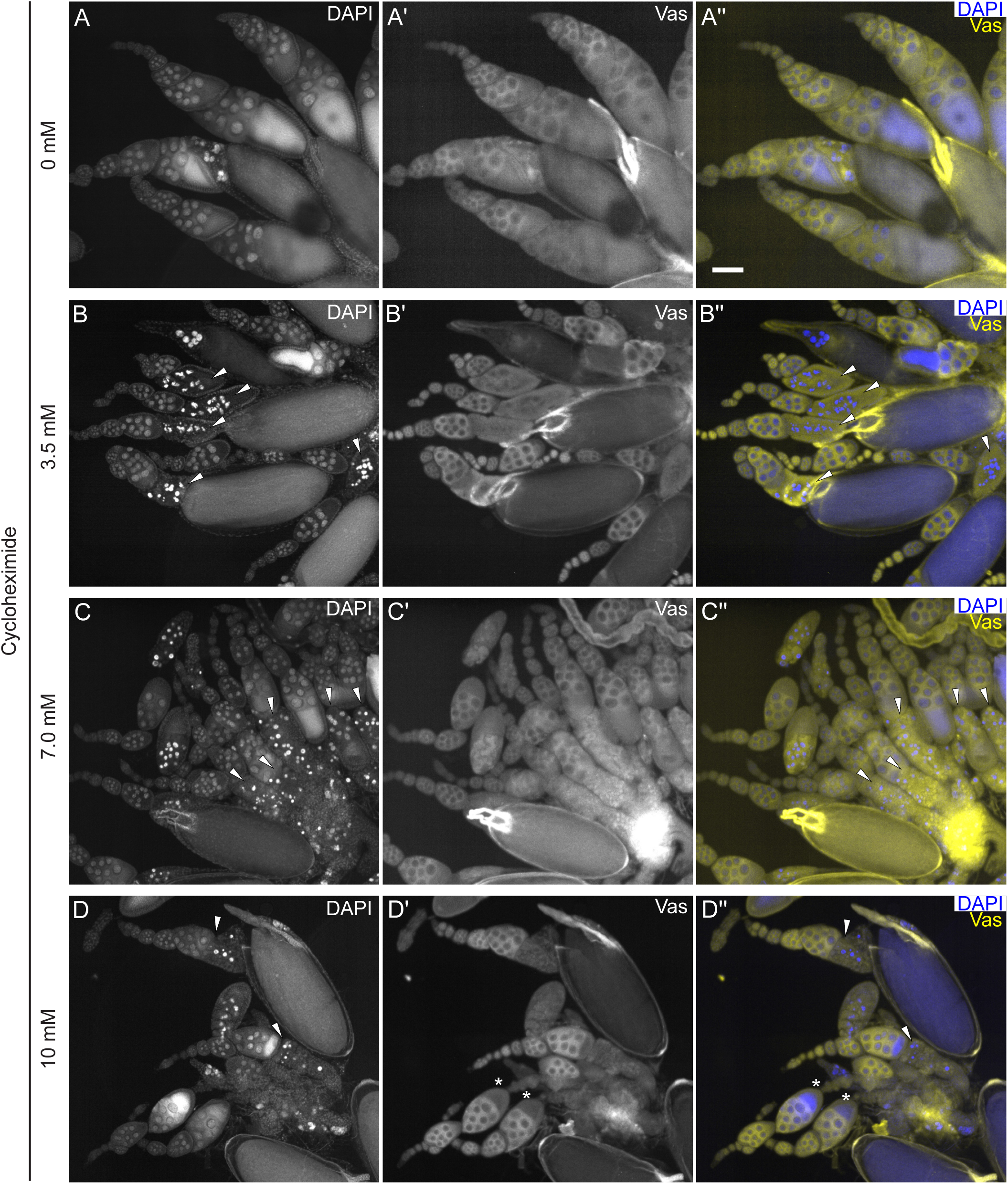
A rich diet supplemented with CHX causes defects in oogenesis. (A-D”) Dissected ovaries labeled with anti-Vasa to label germplasm and DAPI to label nuclei. Ovaries were dissected from females fed for 24 hrs with wet yeast paste containing (A-A”) 0 mM, (B-B”) 3.5 mM, (C-C”) 7.0 mM, and (D-D”) 10 mM CHX. White arrowheads indicate apoptotic cells (B-D”). Asterisks indicates ovarioles with missing follicles stages (D’, D”). (A, B, C, D) White = DAPI. (A’, B’, C’, D’) white = anti-Vasa. (A”, B”, C”, D”, merge) yellow = anti-Vasa, blue = DAPI. Scale bar = 100 µm in A” for A-D”.

**Supplementary Fig 2.**
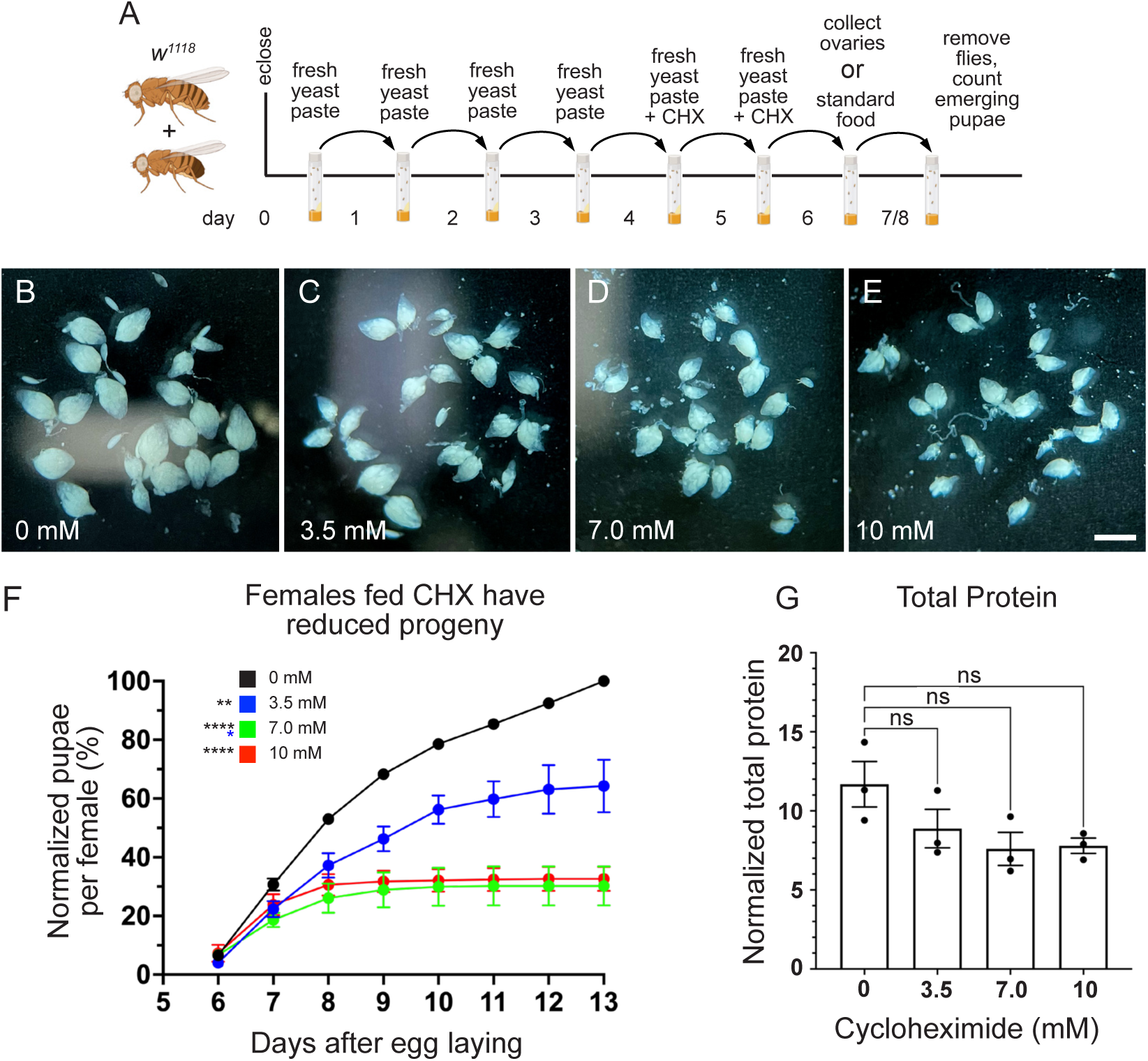
A rich diet supplemented with CHX affects ovary size and progeny number. (A) Schematic illustrating the feeding regimen for comparison of ovary sizes in (B-E) and the pupation assay (F). Newly eclosed flies were fed freshly prepared yeast paste daily for four days, after which they were fed with fresh yeast paste containing CHX for 48 hrs. After CHX feeding, flies were dissected to obtain ovaries or transferred to a standard food vial and allowed to lay eggs for 48 hrs. Subsequently, the adults were removed from the vials and emerging pupae counted. (B-E) Ten pairs of ovaries from females fed with varying concentrations of CHX for 48 hrs. Females fed CHX have visibly smaller ovaries. (F) The percent pupae from females fed with varying concentrations of CHX for 48 hrs normalized to wild type. Percent pupae emerging for the flies fed with CHX-containing yeast paste were significantly lower. (G) Total protein amounts from ovaries after CHX feeding for 24 hrs. The protein amount was normalized with the ovary area (Fig 5F). Each points on the graph represent one replicate, and bars on the graph represent the average of all three replicates. All experimental conditions had three replicates. Error bars = SEM. Data were analyzed for significant differences by ANOVA followed by Tukey’s post hoc test. Asterisks in graphs indicate significance on day 9 after egg laying. * = p < 0.05, ** = p < 0.01, ****= p < 0.0001 and ns= no significance. The color of asterisks in the graphs denotes the concentration of comparison.

## Notes

### Competing Interest Statement

The authors have declared no competing interest.

## References

1. Armstrong, A. R. (2020). Drosophila melanogaster as a model for nutrient regulation of ovarian function. Reproduction, 159(2), R69–R82. 10.1530/REP-18-0593

2. Belozerov, V. E., Ratkovic, S., McNeill, H., Hilliker, A. J., & McDermott, J. C. (2014). In vivo interaction proteomics reveal a novel p38 mitogen-activated protein kinase/Rack1 pathway regulating proteostasis in Drosophila muscle. Mol Cell Biol, 34(3), 474–484. 10.1128/MCB.00824-13

3. Cervantes-Sandoval, I., Chakraborty, M., MacMullen, C., & Davis, R. L. (2016). Scribble Scaffolds a Signalosome for Active Forgetting. Neuron, 90(6), 1230–1242. 10.1016/j.neuron.2016.05.010

4. Darvishi, E., & Woldemichael, G. M. (2016). Cycloheximide Inhibits Actin Cytoskeletal Dynamics by Suppressing Signaling via RhoA. J Cell Biochem, 117(12), 2886–2898. 10.1002/jcb.25601

5. Drummond-Barbosa, D., & Spradling, A. C. (2001). Stem cells and their progeny respond to nutritional changes during Drosophila oogenesis. Dev Biol, 231(1), 265–278. 10.1006/dbio.2000.0135

6. Han, S. K., Lee, D., Lee, H., Kim, D., Son, H. G., Yang, J.-S., Lee, S.-J. V., & Kim, S. (2016). OASIS 2: online application for survival analysis 2 with features for the analysis of maximal lifespan and healthspan in aging research. Oncotarget, 7(35), 56147–56152. 10.18632/oncotarget.11269

7. Jelen, M., Musso, P. Y., Junca, P., & Gordon, M. D. (2023). Optogenetic induction of appetitive and aversive taste memories in Drosophila. Elife, 12. 10.7554/eLife.81535

8. Jeong, E. M., Kwon, M., Cho, E., Lee, S. H., Kim, H., Kim, E. Y., & Kim, J. K. (2022). Systematic modeling-driven experiments identify distinct molecular clockworks underlying hierarchically organized pacemaker neurons. Proc Natl Acad Sci U S A, 119(8). 10.1073/pnas.2113403119

9. Jevtov, I., Zacharogianni, M., van Oorschot, M. M., van Zadelhoff, G., Aguilera-Gomez, A., Vuillez, I., Braakman, I., Hafen, E., Stocker, H., & Rabouille, C. (2015). TORC2 mediates the heat stress response in Drosophila by promoting the formation of stress granules. J Cell Sci, 128(14), 2497–2508. 10.1242/jcs.168724

10. Kim, H. S., Parker, D. J., Hardiman, M. M., Munkacsy, E., Jiang, N., Rogers, A. N., Bai, Y., Brent, C., Mobley, J. A., Austad, S. N., & Pickering, A. M. (2023). Early-adulthood spike in protein translation drives aging via juvenile hormone/germline signaling. Nat Commun, 14(1), 5021. 10.1038/s41467-023-40618-x

11. Lagasse, F., Devaud, J. M., & Mery, F. (2009). A switch from cycloheximide-resistant consolidated memory to cycloheximide-sensitive reconsolidation and extinction in Drosophila. J Neurosci, 29(7), 2225–2230. 10.1523/JNEUROSCI.3789-08.2009

12. Lee, P. T., Lin, G., Lin, W. W., Diao, F., White, B. H., & Bellen, H. J. (2018). A kinase-dependent feedforward loop affects CREBB stability and long term memory formation. Elife, 7. 10.7554/eLife.33007

13. Lin, M. D., Jiao, X., Grima, D., Newbury, S. F., Kiledjian, M., & Chou, T. B. (2008). Drosophila processing bodies in oogenesis. Dev Biol, 322(2), 276–288. 10.1016/j.ydbio.2008.07.033

14. Marcos, R., Lloberas, J., Creus, A., Xamena, N., & Cabre, O. (1982). Effect of cycloheximide on different stages of Drosophila melanogaster. Toxicol Lett, 13(1-2), 105–112. 10.1016/0378-4274(82)90145-x

15. Moutaoufik, M. T., El Fatimy, R., Nassour, H., Gareau, C., Lang, J., Tanguay, R. M., Mazroui, R., & Khandjian, E. W. (2014). UVC-induced stress granules in mammalian cells. PLoS One, 9(11), e112742. 10.1371/journal.pone.0112742

16. Schindelin, J., Arganda-Carreras, I., Frise, E., Kaynig, V., Longair, M., Pietzsch, T., Preibisch, S., Rueden, C., Saalfeld, S., Schmid, B., Tinevez, J.-Y., White, D. J., Hartenstein, V., Eliceiri, K., Tomancak, P., & Cardona, A. (2012). Fiji: an open-source platform for biological-image analysis. Nature Methods, 9(7), 676–682. 10.1038/nmeth.2019

17. Sheth, U., & Parker, R. (2003). Decapping and decay of messenger RNA occur in cytoplasmic processing bodies. Science, 300(5620), 805–808. 10.1126/science.1082320

18. Spradling, A. C. (1993). Developmental genetics of oogenesis. (Cold Spring Harbor Laboratory Press, Cold Spring Harbor)

19. Takakura, M., Nakagawa, R., Ota, T., Kimura, Y., Ng, M. Y., Alia, A. G., Okuno, H., & Hirano, Y. (2021). Rpd3/CoRest-mediated activity-dependent transcription regulates the flexibility in memory updating in Drosophila. Nat Commun, 12(1), 628. 10.1038/s41467-021-20898-x

20. Tully, T., Preat, T., Boynton, S. C., & Del Vecchio, M. (1994). Genetic dissection of consolidated memory in Drosophila. Cell, 79(1), 35–47. 10.1016/0092-8674(94)90398-0

21. Wei, Y., & Lilly, M. A. (2014). The TORC1 inhibitors Nprl2 and Nprl3 mediate an adaptive response to amino-acid starvation in Drosophila. Cell Death Differ, 21(9), 1460–1468. 10.1038/cdd.2014.63

22. Wong, R., Piper, M. D., Wertheim, B., & Partridge, L. (2009). Quantification of food intake in Drosophila. PLoS One, 4(6), e6063. 10.1371/journal.pone.0006063

