## Supplementary material for "Feeding a rich diet supplemented with the translation inhibitor cycloheximide decreases lifespan and ovary size in Drosophila": Supp Figs 1 & 2

Supplemental Fig 1. Cycloheximide feeding causes defects in oogenesis

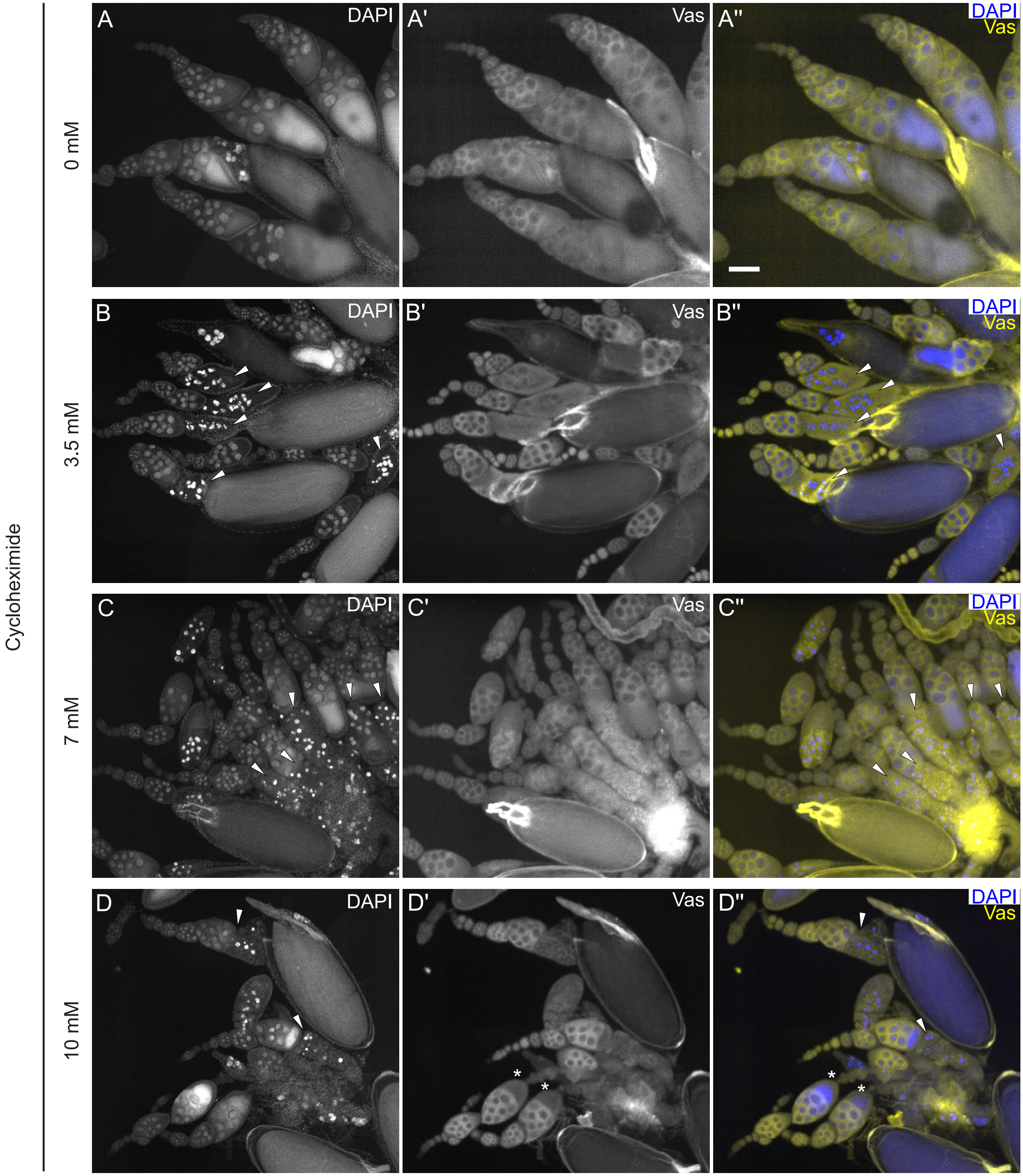

Supplemental Fig 2. Cycloheximide feeding affects ovary sizes and progeny development

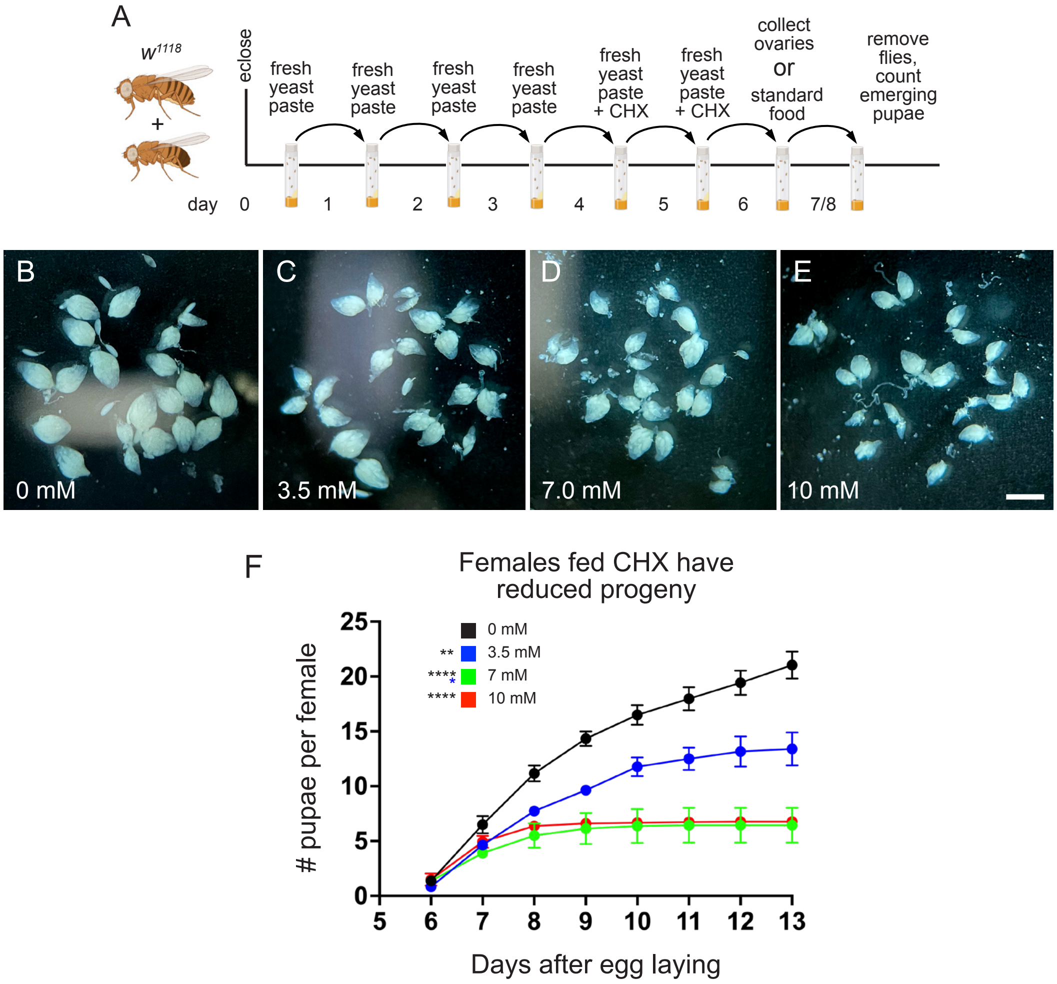
